# Dual targeting of RET and SRC synergizes in RET fusion-positive cancer cells

**DOI:** 10.1101/2025.04.24.650465

**Authors:** Juhyeon Son, Lily L. Remsing Rix, Bin Fang, Eric A. Welsh, Nicole V. Bremer, Valentina Foglizzo, Paola Roa, Nickole Sigcha-Coello, Eric B. Haura, Alexander Drilon, John M. Koomen, Emiliano Cocco, Uwe Rix

## Abstract

*RET* fusions drive subsets of non-small cell lung cancer (NSCLC) and papillary thyroid carcinoma (PTC). Despite new selective RET tyrosine kinase inhibitors (TKIs) resistance usually occurs and is often driven by *RET*-independent bypass mechanisms. Previous studies have implied crosstalk between RET and SRC, but the anti-cancer effects of targeting SRC combined with selective RET TKIs and the underlying molecular mechanisms are not fully understood. Our results showed that the multitargeted SRC TKI dasatinib significantly enhanced efficacy of RET TKIs in *RET* fusion-positive (*RET*^+^) NSCLC and PTC cells. Genetic rescue experiments validated that the combination effects between RET TKIs and dasatinib were indeed SRC-dependent. Phosphoproteomics analysis and validation using selective inhibitors and siRNAs determined that synergy was primarily mediated by suppression of downstream PAK signaling, with contributions from AKT and S6. Importantly, synergy was also observed with eCF506 (NXP900), a next-generation clinical SRC inhibitor. Finally, both SRC TKIs restored sensitivity in selpercatinib-resistant *RET^+^*PTC cells. These results elucidate RET and SRC signaling crosstalk in *RET^+^* NSCLC and PTC suggesting that co-inhibiting SRC has clinical potential in TKI-naïve and -resistant *RET^+^* cancers.

## 1. Introduction

Rearranged during transfection (RET) is a receptor tyrosine kinase (RTK) which is important for the development of normal enteric nervous systems and embryonic kidney, as well as spermatogenesis [1]. *RET* is vulnerable to gene rearrangements driven by inversions of chromosome 10 or translocations with other chromosomes, and those events result in fusion of the *RET* gene with partner genes, e.g., *KIF5B*, *CCDC6* and *NCOA4* [2]. In contrast to wild-type RET, which is strictly regulated by extracellular ligands such as glial cell line-derived neurotrophic factor (GDNF), RET fusions lose regulatory and transmembrane domains and are constitutively activated to promote downstream signaling pathways including mitogen-activated protein kinase (MAPK), PI3K/AKT/mTOR, and JAK2/STAT3 pathways, which are involved in cellular growth, survival and proliferation [1, 3]. *RET* fusions affect 1-2% of non-small cell lung cancer (NSCLC) and 5-10% of papillary thyroid carcinoma (PTC) patients [4–6]. Since *RET* fusion-positive (*RET*^+^) cancers are not very responsive to chemotherapy and immunotherapy, the first line treatment option for patients is targeted therapy using the RET-selective tyrosine kinase inhibitors (TKIs) pralsetinib (BLU-667) or selpercatinib (LOXO-292) [7–9]. Despite the improvement in clinical outcomes by targeted therapy, there are still reports of resistance including acquired *RET* mutations (e.g., V804M gatekeeper and G810R solvent front mutations) and adaptive signals (e.g., via *MET* or *KRAS* amplification) [10–12]. Recently, the next-generation RET TKI vepafestinib (TAS0953/HM06), which has anti-tumor activity against *RET* solvent front mutations both *in vitro* and *in vivo*, has entered a phase 1/2 clinical trial (NCT04683250) [13]. However, concerns remain that many drug resistance mechanisms are *RET*-independent [10]. Thus, discovery and targeting of relevant proteins and mechanisms responsible for resistance are crucial to further improving clinical benefits of RET TKIs by exerting combination effects.

SRC is a non-receptor tyrosine kinase and has been studied for decades as a major target to treat cancer [14]. SRC is involved in essential biological processes such as early embryonic development and immune cell signaling, and its activity in normal cells is tightly regulated by several mechanisms including phosphorylation of inhibitory Y530 by CSK and dephosphorylation of activating Y419 by e.g. phosphatase PTEN [15–17]. Although the *SRC* gene is rarely amplified or mutated in cancers, SRC is frequently overexpressed in tumors and has been reported to contribute to the activation of multiple signaling and scaffold proteins such as RTKs (e.g., EGFR, FGFR, VEGFR, PDGFR and IGF1R) and downstream effectors including MAPKs, FAK, paxillin, p130Cas and RAC1 [14, 18]. Molecular signals driven by SRC result in cell survival, proliferation, migration, stemness and angiogenesis [14]. Due to its involvement in several cellular functions and its ability to contribute to the activation of driver oncogenes, SRC has been proposed to facilitate tumor adaptation to therapeutic stress by promoting the activation of compensatory pathways.

Consistently, previous studies showed that inhibition of SRC can strengthen the efficacy of targeted therapies to treat NSCLCs harboring driver oncogenes such as mutant *EGFR*, *ALK* fusions, and *ROS1* fusions [19–22]. This preclinical evidence is further supported by clinical studies using the EGFR inhibitor erlotinib and the SRC inhibitor dasatinib to treat patients with advanced NSCLC [23, 24]. More specific to *RET*^+^ cancer, a previous study found that *CCDC6-RET-*driven LC-2/Ad NSCLC cells with acquired resistance to the unselective FGFR, VEGFR, and PDGFR inhibitor, dovitinib, which also has non-canonical activity against RET, were sensitive to SRC inhibition [25]. In addition, various thyroid cancer cell lines including *CCDC6-RET-*driven TPC-1 PTC cells were sensitive to single agent dasatinib resulting in their growth suppression [26]. Other investigations reported that KIF5B-RET fusions interact with SRC to induce invadopodia-like processes including cell migration and invasion in *Drosophila* cells [27]. Taken together, it is implied that SRC is a potential candidate accountable for the bypass mechanisms in *RET*^+^ cancers. However, it is still not clearly understood whether targeting SRC sensitizes the *RET*^+^ cancer cells to the selective RET TKIs selpercatinib and pralsetinib, which are current first line therapies. Furthermore, mechanistic studies are necessary to identify and validate the downstream signaling proteins and pathways targeted by co-treatment with RET and SRC inhibitors in *RET*^+^ positive cancers. Thus, we investigated the drug combination effects of co-targeting RET and SRC to enhance drug response in *RET*^+^ cancer cells and the related molecular mechanisms by conducting phosphoproteomics and functional validation assays.

## 2. Materials and Methods

### 2.1. Cell lines and reagents

CUTO32 and LC-2/Ad cell lines were obtained from Dr. Robert Doebele at the University of Colorado as described in a previous report [28], while the TPC-1 cell line was obtained by Dr. Fagin at the Memorial Sloan Kettering Cancer Center [29]. All cell lines were tested negative for mycoplasma and LC-/Ad cells were authenticated with STR analysis. Cells were cultured in RPMI1640 (Corning, MT10040CV) media containing 10% of fetal bovine serum (Sigma, F2442) and humidified atmosphere with 5% CO_2_. Pralsetinib, selpercatinib, PF-3758309, and eCF506 were obtained from MedChemExpress. GSK690693 was purchased from Selleckchem. Dasatinib was obtained from ChemieTek and rapamycin from LC Laboratories. All compounds were diluted with dimethylsulfoxide (DMSO, Acros Organics) to a final concentration of 10 mM and stored at −20°C. For drug treatment experiments, DMSO was used as a vehicle control.

### 2.2. Cell Viability

Cells were plated in 384-well black plates with clear bottom with a density of 1,000 cells per well or in 96-well plates. After 24 hours, drugs diluted in media at the indicated concentrations were added to each well. After 3 days of drug treatment, CellTiter-Glo (Promega) reagent was added according to the manufacturer’s instructions. Luminescence which indicates cell viability was measured by Spectramax M5 plate reader (Molecular Devices) connected with SoftMax Pro 7.1 at 500 ms integration time. The synergistic effect was evaluated using Bliss independence model. ΔBliss score greater than 0.05 was considered synergistic, less than −0.05 as antagonistic, and between −0.05 and 0.05 as indicative of an additive effect.

### 2.3. Clonogenicity

CUTO32 cells were plated at a density of 1 x 10^5^ cells in 6-well plates or 4 x 10^4^ cells in 12-well plates. LC-2/Ad cells were plated at a density of 2 x 10^4^ cells in 6-well plates or 1 x 10^4^ cells in 12-well plates. TPC-1 were seeded at a density of 4 x 10^3^ cells in a 24-well plate. Cells were treated with drugs at the indicated concentrations for 5 to 7 days, replacing with fresh media every 3-4 days. At the end point of treatment, cells were washed with phosphate-buffered saline (PBS, D8537, Sigma) once, fixed for 30 min on ice with methanol, and stained with crystal violet solution (Sigma, HT90132) for 30 min at room temperature (RT). Crystal violet was extracted using 2 mL of methanol and transferred to 96-well plate with clear bottom. Absorbance of extracted crystal violet was measured by Spectramax M5 plate reader at 540 nm wavelength. The synergistic effect was evaluated using the Bliss independence model.

### 2.4. Western blotting

Cells were harvested with ice-cold PBS containing sodium vanadate, and lysed with a lysis buffer (100 mM NaCl, 1.5 mM MgCl_2_, 5% glycerol, 0.2% NP-40, 50 mM Tris, pH7.5) mixed with phosphatase inhibitor cocktail 2 (Sigma, P5726) and cOmplete protease inhibitor cocktail (Roche, 11873580001). Extracted proteins were separated by SDS-PAGE and transferred to PVDF membranes using Trans-Blot Turbo Transfer System (BioRad, 1704150EDU). Membranes were blocked with aqueous 5% non-fat dry milk in TBS-T buffer and incubated with primary antibodies overnight at 4°C. Next day, membranes were incubated with horseradish peroxidase (HRP)-conjugated secondary antibodies for 1 hour at room temperature. Clarity Western ECL Substrate (BioRad, 1705061) or Clarity Max Western ECL Substrate (BioRad, 1705062) were added to the membranes and images were visualized and quantified by Odyssey Fc LI-COR Dual-Mode Imaging System and LI-COR’s Image Studio Lite (version 5.2) software. Primary antibodies against PARP1 (9542), cleaved Caspase-3 (9661), p-RET (Y905, 3221), RET (3223) p-SRC (Y419, 2101), SRC (2108), p-PAK1/2 (S144/S141, 2606), PAK1/2/3 (2604), p-ERK1/2 (Y204/Y187, 4370), p-AKT (S473, 9271), AKT (9272), p-S6 (S235/S236, 4858), S6 (2217) and Vinculin (13901) were acquired from Cell Signaling Technology. Primary antibodies against ERK1/2 (M5670) and β-Actin (A5441) were obtained from Sigma. Anti-rabbit (45000682) and anti-mouse (45000679) secondary antibodies were purchased from Fisher Scientific.

### 2.5. Phospho- and expression proteomics

CUTO32 cells (1.8 x 10^6^ cells per 100 mm culture dish, 3 biological replicates) were treated with DMSO or drugs at indicated concentrations for 3 hours. Drug-treated cells were harvested with ice-cold PBS and shock frozen with LN_2_. Cells were lysed in denaturing lysis buffer containing aqueous 8 M urea, 20 mM HEPES (pH 8), 1 mM sodium orthovanadate, 2.5 mM sodium pyrophosphate and 1 mM β-glycerophosphate. Bradford assays were carried out to determine the protein concentration in each sample. The disulfide bonds were reduced with 4.5 mM dithiothreitol (DTT), and the resulting cysteines were alkylated with 10 mM iodoacetamide. Trypsin digestion was carried out at room temperature overnight, and tryptic peptides were then acidified with aqueous 1% trifluoroacetic acid (TFA) and desalted with C18 Sep-Pak cartridges according to the manufacturer’s procedure. Peptides from each sample were labeled with TMTPro 18plex reagent to barcode them to sample of origin. Label incorporation was checked by LC-MS/MS and spectral counting for TMT-labeled peptides; quality control passed when 95% or greater label incorporation was achieved for each channel. The 18 samples were then pooled using equal amounts of total protein digest and lyophilized. After lyophilization, the peptides were re-dissolved in 400 μL of aqueous 20 mM ammonium formate, (pH 10.0), basic pH reversed phase liquid chromatography (bRPLC) solvent A. The bRPLC separation was performed on an XBridge 4.6 mm ID x 100 mm length column packed with BEH C18 resin, 3.5 µm particle size, 130 Å pore size. (Waters) The peptides were eluted as follows: 5% bRPLC solvent B (aqueous 5 mM ammonium formate, 90% acetonitrile, pH 10.0) for 10 minutes, 5% - 15% B in 5 minutes, 15-40% B in 47 minutes, 40-100% B in 5 minutes and 100% B held for 10 minutes, followed by re-equilibration at 1% B.

The flow rate was 0.6 mL/min, and 12 concatenated fractions were collected for phosphopeptide enrichment; 24 concatenated fractions were collected for protein expression. Vacuum centrifugation (Speedvac, Thermo) was used to dry the peptide fractions. Peptides were re-dissolved in aqueous IMAC loading buffer containing 0.1% TFA and 85% acetonitrile (ACN). The phosphopeptides in each fraction were enriched using IMAC resin (#20432, Cell Signaling Technology) using a KingFisher 96 (ThermoFisher). Briefly, the IMAC resin was washed once with loading buffer. The peptides were incubated with the IMAC resin for 30 minutes at room temperature with gentle agitation. Ten microliters magnetic bead IMAC resin was added per sample. After incubation, the IMAC resin was washed twice with loading buffer followed by 1 wash with wash buffer (aqueous 80% ACN containing 0.1% TFA). The phosphopeptides were eluted with elution buffer (aqueous 50% ACN containing 2.5% ammonia). The volume was reduced to 20 µL via vacuum centrifugation. A nanoflow ultra-high performance liquid chromatograph (RSLCnano, Dionex, Sunnyvale, CA) interfaced with an electrospray orbitrap mass spectrometer (Orbitrap Exploris 480, Thermo, San Jose, CA) was used for tandem mass spectrometry peptide sequencing experiments. The sample was first loaded onto a pre-column (C18 PepMap100, 100 µm ID x 2 cm length packed with C18 reversed-phase resin, 5 µm particle size, 100 Å pore size) and washed for 8 minutes with aqueous 2% ACN and 0.04% TFA. The trapped peptides were eluted onto the analytical column, (C18 PepMap100, 75 µm ID x 25 cm length, 2 µm particle size, 100 Å pore size, Thermo). The 120-minute gradient was programmed as: 95% solvent A (aqueous 2% ACN + 0.1% formic acid) for 8 minutes, solvent B (aqueous 90% ACN + 0.1% formic acid) from 5% to 38.5% in 90 minutes, then solvent B from 50% to 90% B in 7 minutes and held at 90% for 5 minutes, followed by solvent B from 90% to 5% in 1 minute and re-equilibration for 10 minutes. The flow rate on analytical column was 300 nL/min. Cycle time was set at 3 sec for data dependent acquisition. Spray voltage was 2,100 V, and capillary temperature was 300 °C. The resolution for MS and MS/MS scans were set at 120,000 and 45,000 respectively. Dynamic exclusion was 15 seconds to prevent additional sequencing of previously sampled peptide peaks. MaxQuant (version 1.6.14.0) was used to identify peptides and quantify the TMT reporter ion intensities [30]. The UniProt database was downloaded March 2023. Up to 2 missed trypsin cleavages were allowed. Carbamidomethyl cysteine was set as fixed modification. Phosphorylation on Serine/Threonine/Tyrosine and oxidation of Methionine were set as variable modifications. Both peptide spectral matching (PSM) and protein false discovery rate (FDR) were set at 0.05. The match between runs feature was activated to enable quantification across samples without requiring MS/MS identification in every analysis.

### 2.6. Processing and analysis of proteomic data

Samples within each multiplex were normalized with IRON (iron_generic --proteomics) against their respective median sample channels (total protein: 130C, pSTY run1: 130C, pSTY run2: 131C), selected as the channel least-different from other channels (findmedian--spreadsheet --pearson) [31]. In order to remove systematic differences in signal between pSTY multiplex replicates, normalized abundances within each pSTY multiplex were converted to ratios against their respective all-channel computational pools (geometric mean abundance, per row, across all channels). The computational pools were normalized together using IRON against their median sample. For each row of data, the geometric mean of the normalized computational pool abundances was calculated and stored. Sample vs. computational pool ratios were then scaled back into abundance values using the stored pool row means. All abundances were then log_2_ transformed prior to additional analyses. Injection replicates (pSTY: n=2, total protein n=1) for each sample were averaged together, then average log2 abundances calculated from the biological replicates within each condition. Log_2_ ratios between conditions were calculated by subtracting average log_2_ abundances. Two-group p-values were calculated using two-sided, unequal-variance Welch’s T-tests from the individual averaged injection replicates within each condition. Scores were calculated as the geometric mean of the |log_2_ ratio| and -log_10_(p-value), multiplied by the sign of the log_2_ ratio. Rows were sorted on number of differentially expressed conditions, |Score|, and direction of change to bring the rows with the strongest potential biological signal to the top of the analysis spreadsheet. For pSTY dataset, phosphosites were filtered by cutoffs with |log_2_ ratio| > 1 [log_2_(2-fold)] and p-value < 0.05 to identify differentially regulated phosphosites. For expression dataset, proteins were filtered by cutoffs with |log_2_ ratio| > 0.585 [log_2_(1.5-fold)] and p-value < 0.05 to find out proteins of which expression levels were significantly affected. All proteins which passed cutoffs were selected, and pathway enrichment analysis was conducted by Database for Annotation, Visualization and Integrated Discovery (DAVID) and Kyoto Encyclopedia of Genes and Genomes (KEGG) database [32, 33]. Enriched pathways which have Benjamini-adjusted p-value less than 0.05 were selected for further analysis. Pathway map from Fig. 3C was created with Biorender.com based on the relevant KEGG pathway maps (hsa04810 and hsa04150). For the network analysis, STRING (version 12.0) was used to analyze protein-protein interactions, and minimum required interaction score was set as low confidence (0.15) [34]. Protein networks were imported to and visualized by Cytoscape (version 3.10.2) and GLay was used for generating network clusters [35, 36]. To identify targets with high priority, betweenness centrality and fold change of protein phosphorylation or expression was plotted using Excel and GraphPad Prism 10.

### 2.7. RNA interference

ON-TARGETplus Human PAK1 siRNA (L-003521-00-0005) and ON-TARGETplus non-targeting (D-001810-10-20) were obtained from Dharmacon. Reverse transfection was conducted according to the manufactuer’s protocol. Briefly, each siRNA (20 nM as a final concentration) was mixed with Lipofectamine™ RNAiMAX Transfection Reagent (ThermoFisher) in Opti-MEM (ThermoFisher). siRNA-Lipofectamine complexes were added to each well of a 6-well plate, followed by the addition of 1 x 10^5^ cells in resuspension. After 24 hours, cells were treated with DMSO or pralsetinib at the indicated concentrations for additional 3 days. Treated cells were harvested and 10% of cell resuspension was used for CellTiter-Glo assay. Rest of cell resuspension was used for western blot to validate knockdown.

### 2.8. Viral transductions

pLenti6/UbC/V5-DEST Gateway Vector plasmids (Invitrogen, V49910) constructed with open reading frames of wild type and gatekeeper mutant SRC were kindly gifted by Drs. Jinyan Du and Todd R. Golub [37]. Lentiviruses were generated by transfecting HEK293T cells with 3^rd^ Generation Packaging System Mix (Applied Biological Materials, LV053), construct plasmids and jetPRIME (VWR, 89129-922) transfection reagent. Media containing viruses were collected and filtered with Millex-HV Filter Unit (Millipore, SLHV033RS). CUTO32 and LC-2/Ad cells were plated in 6-well plates (1.5 x 10^5^ cells per well) and transduced with collected viruses and 8 μg/mL of polybrene (Millipore, TR-1003-G). After 48 hours, transduced cells were selected by adding 10 μg/mL of blasticidin (ThermoFisher, J67216.XF) for 7 days.

### 2.9. Quantification and Statistical analysis

Raw data were processed and analyzed by Excel and GraphPad Prism 10. Data presented in graphs are mean ± standard deviation, defining the values of DMSO-treated cells as 100% or 1-fold change. Statistical significance was determined by Welch’s t-test or one-way ANOVA with Tukey’s multiple comparisons, using p-value less than 0.05 to determine statistical significance.

## 3. Results

3.1. Dasatinib synergizes with RET TKIs in *RET* fusion-positive cancer cells

To determine whether the dual targeting of RET and SRC exerts synergy to treat RET^+^ cancer, we treated CUTO32 (*KIF5B-RET*), LC-2/Ad (*CCDC6-RET*) NSCLC and TPC-1 PTC cell lines with a combination of RET TKIs (pralsetinib or selpercatinib) and dasatinib. In all three cell lines, combination with RET TKIs and dasatinib exerted synergistic effects compared to single drug treatment (Fig. 1A and Supplementary Fig. S1A). Those results were also consistent with observations from long term clonogenicity assays (Fig. 1B and Supplementary Fig. S1B). To analyze apoptosis induced by drug combinations, we performed western blotting and detected cleaved PARP1 and cleaved caspase-3, which are major apoptosis markers. While single agent dasatinib did not cause cell death compared to negative control in CUTO32 cells, it further promoted apoptosis induction when combined with pralsetinib compared to pralsetinib alone (Fig. 1C). In summary, our results suggest that SRC inhibition may improve sensitivity of naïve *RET*^+^ cancer cells to RET-targeted therapy.

**Figure 1.**
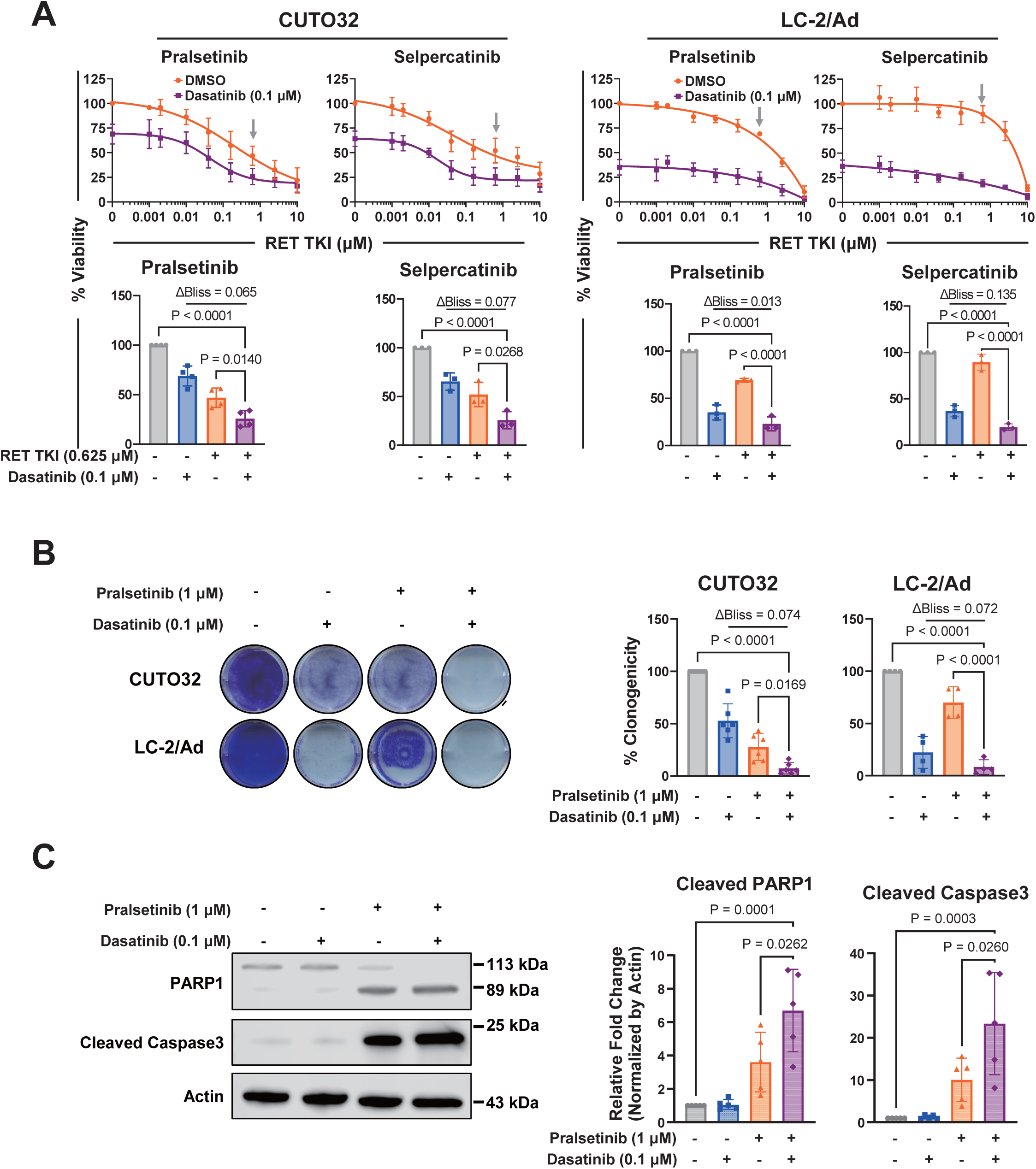
Dasatinib enhancement of RET TKI efficacy in *RET^+^* NSCLC cells. (A) Cell viability of *RET*^+^ CUTO32 and LC-2/Ad NSCLC cells. (Top) Dose response curves of RET TKIs (pralsetinib, selpercatinib) combined with dasatinib (0.1 μM) for 3 days. (Bottom) Cell viability and Bliss synergy analysis of *RET*^+^ NSCLC cells at the single concentration of RET TKIs (0.625 μM, see grey arrows above). (B) Clonogenic survival, quantification and Bliss synergy analysis of *RET*^+^ NSCLC cells after co-treatment with pralsetinib (1 μM) and dasatinib (0.1 μM) for 7 days. (C) Western blotting of apoptosis markers cleaved PARP1 and cleaved caspase-3 after co-treatment with pralsetinib (1 μM) and dasatinib (0.1 μM) for 48 hours in CUTO32 cells.

### 3.2. Synergistic effects of dasatinib with RET TKIs are SRC-dependent

Although dasatinib is a well-known compound to strongly inhibit SRC, this drug is reported to have a broad range of off-targets including other SRC family kinases (e.g., LYN, YES and LCK), ABL, TEC, GAK, and p38α [38]. Since this drug profile of dasatinib entails the possibility of polypharmacology in terms of treating *RET*^+^ cancer cells, it is challenging to conclude that the synergy between RET TKIs and dasatinib is SRC-dependent. To validate whether the effects of dasatinib in *RET*^+^ NSCLC cells were driven indeed by SRC inhibition, the cells were infected with lentivirus carrying wild-type SRC or the gatekeeper-mutant SRC (T341I), which prevents dasatinib binding (Fig. 2A) [37, 38]. As expected, immunoblotting for SRC autophosphorylation indicated that the activity of gatekeeper SRC in both CUTO32 and LC-2/Ad cells was not suppressed by dasatinib treatment compared to wild-type SRC (Fig. 2B and Supplementary Fig. S2A). Importantly, this difference was also reflected by the results from viability and clonogenicity assays, which showed much weaker effects of the combination treatment with RET TKIs and dasatinib in both cell lines expressing the SRC gatekeeper mutant compared to the cells expressing wild-type SRC (Fig. 2C, D and Supplementary Fig. S2B, C). These results indicate that synergy of dasatinib with RET TKIs in *RET*+ NSCLC cells is mediated by inhibition of its canonical target SRC with little to no contribution from dasatinib off-targets.

**Figure 2.**
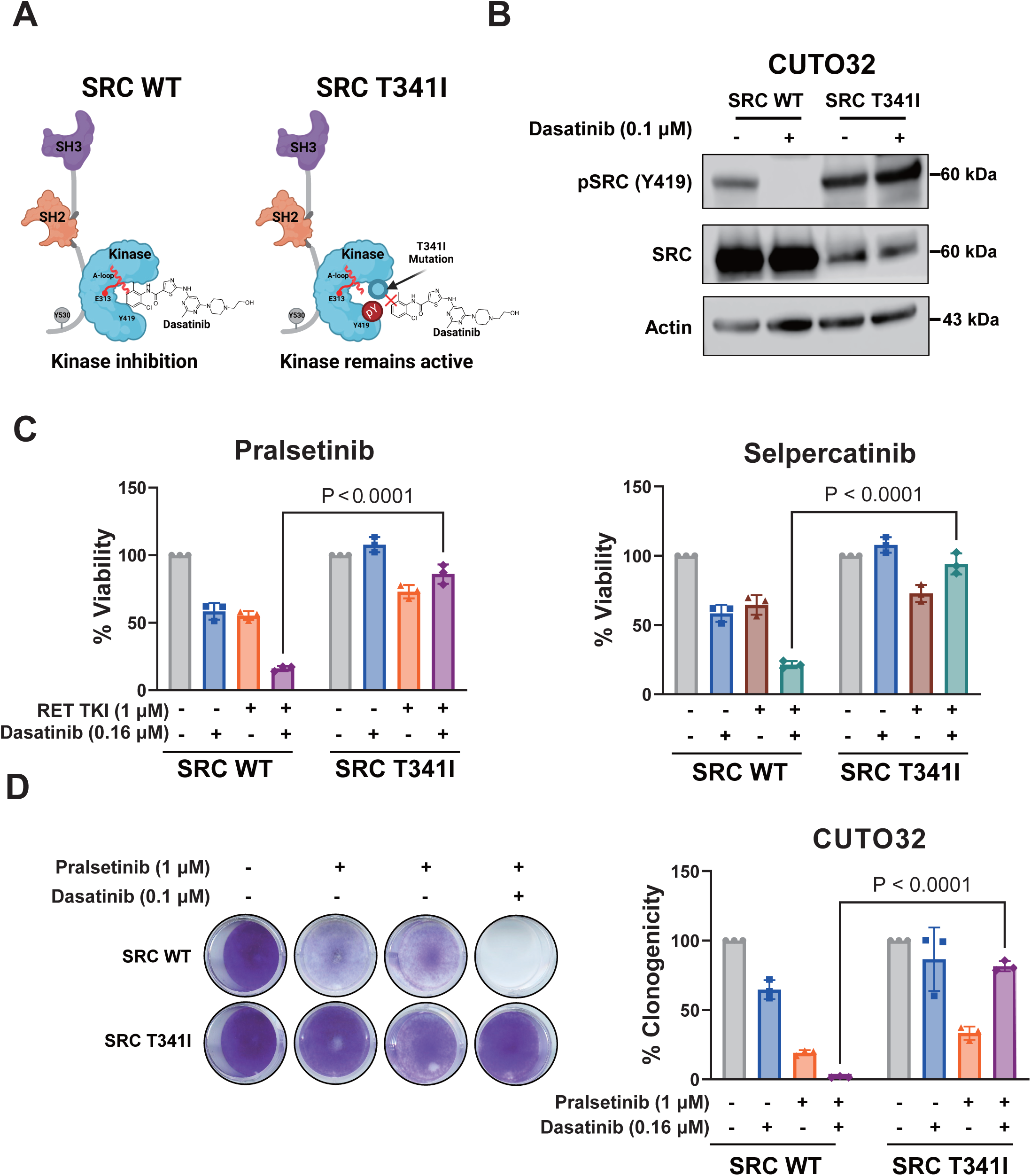
SRC-dependency of synergistic effects between RET TKIs and dasatinib in *RET^+^* CUTO32 NSCLC cells. (A) Schematic representation of the mechanism through which the SRC gatekeeper mutation (T341I) disrupts dasatinib binding. Scheme was adapted from a study by Temps et al [44]. (B) SRC autophosphorylation in CUTO32 cells expressing SRC wild-type (WT) or T341I SRC gatekeeper mutation upon treatment with dasatinib (0.1 μM) for 3 hours. (C) Cell viability of CUTO32 cells expressing SRC WT or SRC T341I gatekeeper mutation after combined treatment with dasatinib (0.16 μM) and the RET TKIs pralsetinib and selpercatinib (both 1 μM) for 3 days. (D) Clonogenic survival and quantification of CUTO32 cells expressing SRC WT or SRC T341I gatekeeper mutation upon treatment with pralsetinib (1 μM), dasatinib (0.1 μM) or combination thereof for 7 days .

**Figure 3.**
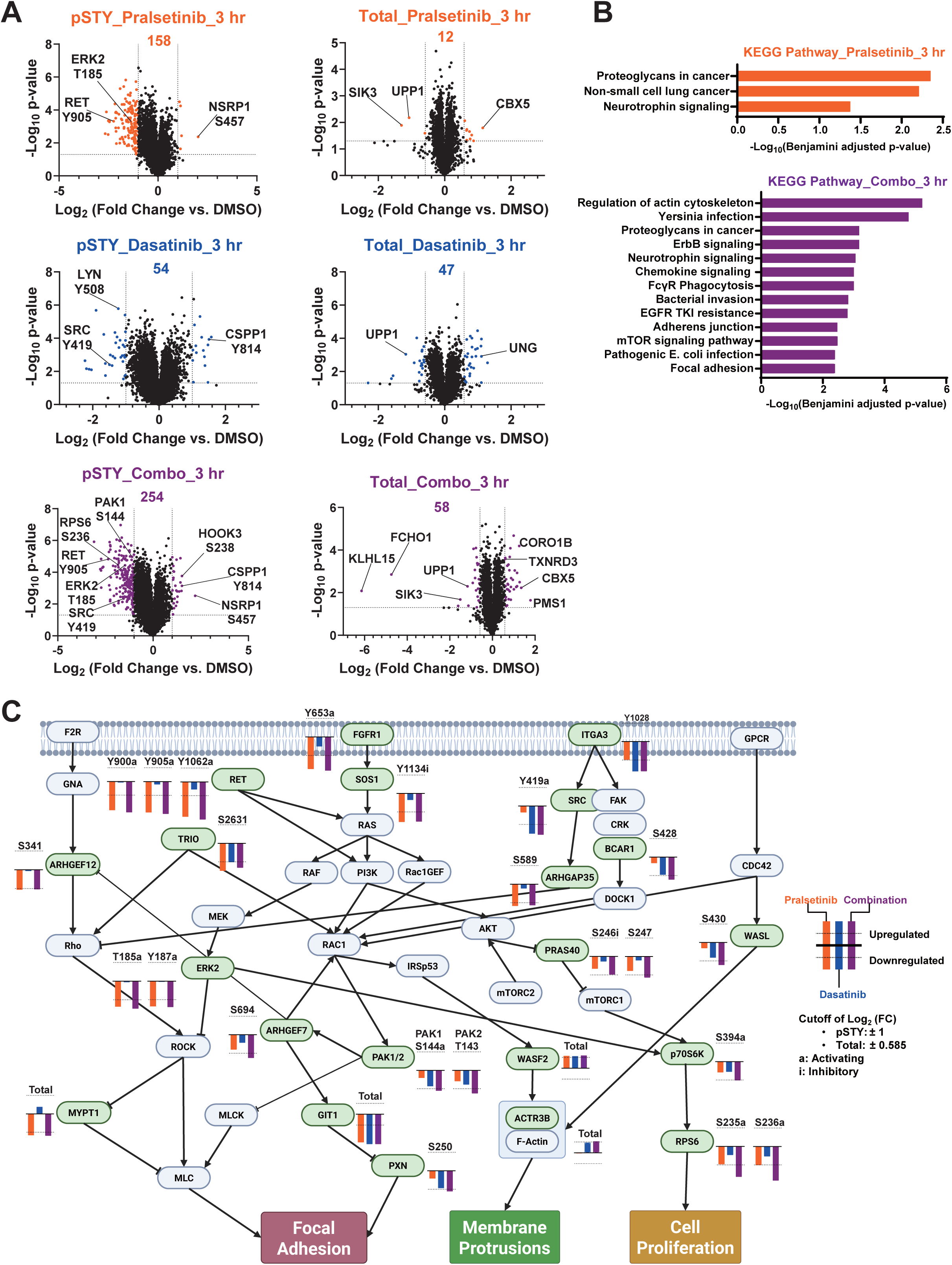
Proteomics analysis of signal changes by treatment with pralsetinib and dasatinib. (A) Volcano plots of proteomics analysis presenting significantly regulated phosphosites (pSTY) and protein expression (Total) upon treatment with pralsetinib (1 μM), dasatinib (0.1 μM) or their combination (Combo) for 3 hours. (B) DAVID analysis of KEGG pathways significantly regulated by pralsetinib or combination treatment. Pathways with Benjamini adjusted p-value less than 0.05 are displayed. (C) Signaling proteins significantly modulated by drug treatment mapped on the coalesced enriched pathways with a plot indicating Log_2_(Fold Change) of phosphorylations or total expression upon each individual treatment. a: activating phosphorylation; i: inhibitory phosphorylation.

### 3.3. Proteomic analysis of signaling changes in RET^+^ NSCLC by combined treatment with pralsetinib and dasatinib

To elucidate how SRC inhibition cooperates with RET inhibitors to suppress RET^+^ cancer cell survival, we aimed to identify proteins, which are downstream of SRC and responsible for adaptive signaling to RET inhibition. For this purpose, we conducted quantitative global (pSTY) phosphoproteomics and expression proteomics analyses after treating CUTO32 cells with pralsetinib, dasatinib, or combination thereof for 3 hours. We detected close to 20,000 phosphosites from global phosphoproteomics data and over 6,000 proteins from the expression proteomics data. Using all phosphopeptides or expressed proteins, which have passed cutoffs of |Fold Change| > 2 (pSTY) or |Fold Change| > 1.5 (expression) and p-value less than 0.05 (from both proteomics datasets), combination treatment significantly modulated 254 phosphosites and 58 proteins while single agent pralsetinib significantly modulated 158 phosphosites and 12 proteins (Fig. 3A). Using all proteins, which have passed the specified cutoffs, we generated a protein network to query and visualize functional protein-protein interactions using STRING (Supplementary Fig. S3A). After plotting those proteins based on the log_2_ ratio of change in phosphorylation or expression levels and their betweenness centrality within the network, we have identified ERK2, PAK1/2, PRAS40 and S6 as high-priority targets (Supplementary Fig. S3B). Also, we conducted pathway enrichment analyses using DAVID database [39]. Compared to single pralsetinib treatment, which enriched only three pathways, particularly MAPK signaling, that collectively represented RTK signaling in NSCLC, combination treatment additionally enriched various other pathways (Fig. 3B), most prominently, regulation of actin cytoskeleton, ERBB, and mTOR signaling pathways. Since the combination treatment did not significantly affect phosphorylation of any ERBB family receptors directly, we focused on regulation of actin cytoskeleton and mTOR signaling pathways as signals potentially responsible for eliciting synergy between pralsetinib and dasatinib. Aggregating the KEGG pathway maps into one, we identified ERK2, PAK1, PRAS40, and S6 as potentially relevant downstream targets, consistent with the protein network analysis (Fig. 3C). In particular, the activating phosphorylations of PAK1 (S141) and S6 (S235/S236) were further decreased by combination treatment compared to single pralsetinib treatment. PAKs are non-receptor serine/threonine kinases downstream of RAC1 with biological functions in proliferation, metastasis, angiogenesis and drug resistance [40]. Although PAKs do not directly interact with SRC, they can be indirectly affected as part of the SRC/VAV/RAC1/PAK signaling axis [41]. It is also notable that the inhibitory site of PRAS40 (T246) was downregulated stronger by drug combination, although its fold change (Log_2_ ratio of −0.97) narrowly missed the cutoff. As this protein is directly inhibited by AKT and negatively regulates mTOR complex 1 (mTORC1) upstream of S6, these results imply that co-targeting RET and SRC further inhibits AKT resulting in stronger suppression of downstream mTORC1 and S6 signals, which are critical for processes including mRNA translation, cell survival and proliferation [42]. Taken together, integrated proteomics analysis suggested that while RET targeting predominantly strongly reduced MAPK signaling, SRC TKI combination markedly enhanced inhibition of PAK and AKT/mTOR signaling.

### 3.4. SRC targeting enhances inhibition of PAK and AKT signaling by pralsetinib

To validate the functional proteomics data, we analyzed whether the combination of pralsetinib and dasatinib affects PAK, AKT and S6 phosphorylation also by immunoblot analysis. CUTO32 cells were treated with pralsetinib, dasatinib, and their combination for 3 hours and western blotting was conducted for phosphorylation of RET, SRC, PAK, ERK, AKT, and S6. As expected, pralsetinib abrogated RET autophosphorylation, which resulted in strong inhibition of pERK. Dasatinib potently inhibited SRC autophosphorylation, but as a single agent predominantly reduced activity-associated phosphorylation of PAK, and to some extent S6 phosphorylation, without significantly suppressing ERK and AKT. It is possible that inhibitory effects against these proteins could be sufficiently compensated by RET. However, combination treatment more strongly suppressed phosphorylation of PAK, AKT and S6 compared to single agent pralsetinib (Fig. 4A). To determine, if these combined signaling effects were attributable to on-target SRC inhibition by dasatinib rather than an off-target, we examined the effects of dasatinib and combination treatment on these signaling pathways in CUTO32 cells expressing either wild-type SRC or the dasatinib-resistant SRC gatekeeper mutant. As observed in parental CUTO32 cells, activating phosphorylation of PAK, AKT and S6 were further reduced by combination treatment in CUTO32 cells expressing wild-type SRC. In contrast, dasatinib-induced inhibition of these signals were completely rescued in cells expressing gatekeeper SRC (Fig. 4B). Overall, while pralsetinib alone predominantly inhibited pERK as the most prominent RET downstream signal, additional targeting of SRC markedly improved blockade of PAK and AKT, leading also to further attenuated activity of downstream S6.

**Figure 4.**
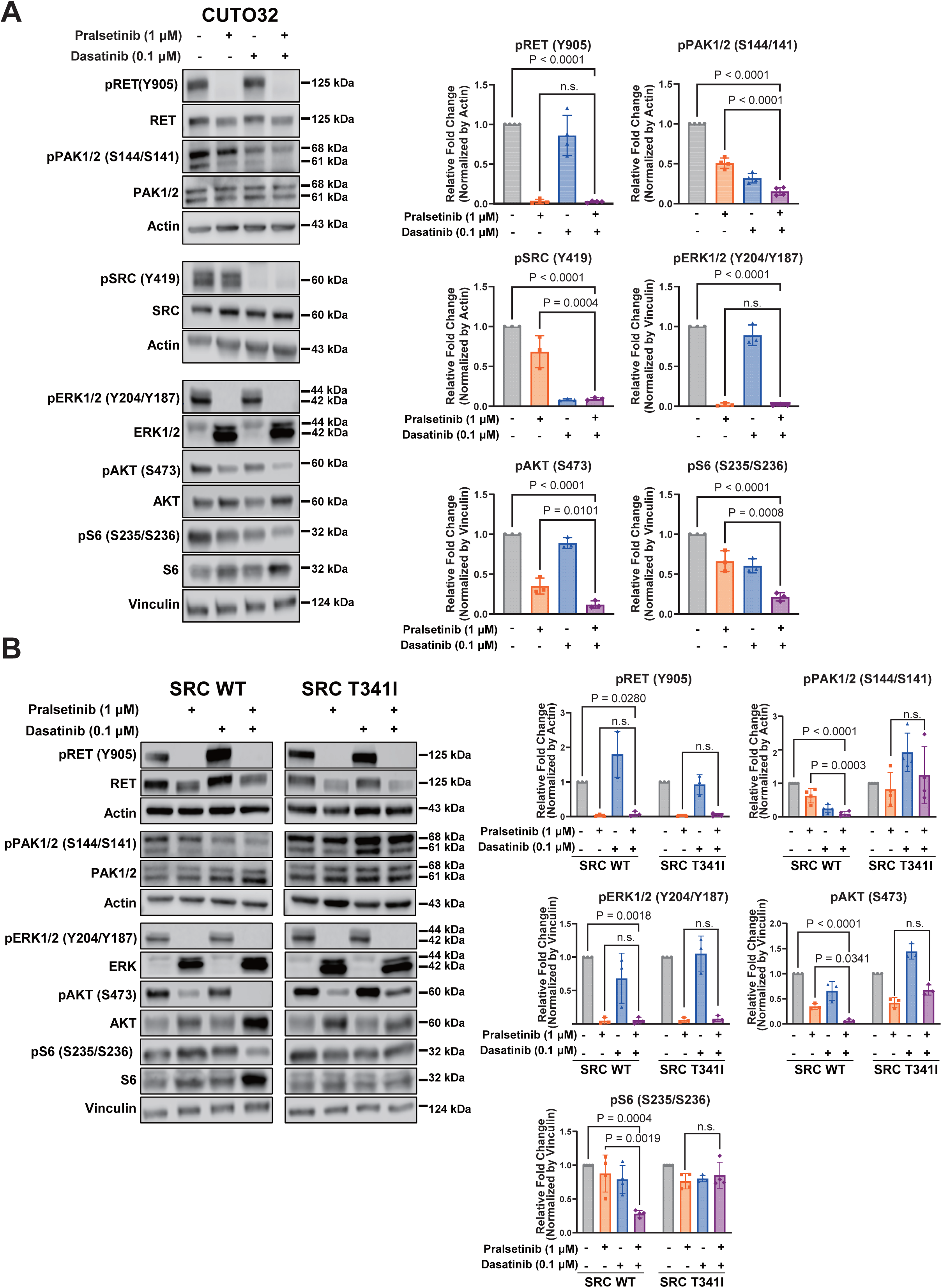
Signaling analysis of pralsetinib and dasatinib combination effects in *RET*^+^ NSCLC cells. (A) (Left) Western blot analysis of indicated signaling proteins in CUTO32 cells after treatment with pralsetinib (1 μM), dasatinib (0.1 μM) or their combination for 3 hours. (Right) Quantification of pRET, pSRC, pPAK1/2, pERK1/2, pAKT, and pS6 blots, normalized by loading controls (actin or vinculin). (B) (Left) Western blot analysis of indicated signaling proteins after treatment with pralsetinib (1 μM), dasatinib (0.1 μM) or their combination for 3 hours in CUTO32 cells expressing SRC wild-type (WT) or SRC T341I gatekeeper. (Right) Quantification of pRET, pPAK1/2, pERK1/2, pAKT, and pS6 blots, normalized by loading controls (actin or vinculin).

### 3.5. Synergy is mediated by cooperative inhibition of PAK and AKT/mTOR signaling

To determine whether inhibition of PAK, AKT and S6 phosphorylation impacts viability of *RET*^+^ NSCLC cells, we treated CUTO32 and LC-2/Ad cells with pralsetinib in combination with the selective AKT inhibitor GSK690693, mTORC1 inhibitor rapamycin or PAK inhibitor PF-3758309. In CUTO32 cells, all three inhibitors enhanced cellular activity of pralsetinib with the effect of PF-3758309 being the strongest (Fig. 5A and Supplementary Fig. S4A). In LC-2/Ad cells, only PF-3758309 could significantly reduce cell viability compared to single pralsetinib treatment (Fig. 5A and Supplementary Fig. S4A). A previous study suggested that PAK1 activates AKT1 via its non-catalytic scaffold function in fibroblasts [43]. To check whether pharmacological PAK inhibition affects AKT signaling in *RET*^+^ NSCLC cells, we treated CUTO32 cells with up to 5 μM PF-3758309 for 3 hours. However, PAK inhibition did not decrease p-AKT and p-S6 in these cells, indicating that the observed dasatinib effects on AKT downstream of SRC are independent from PAK inhibition in these cells (Supplementary Fig. S4B). Since PAK inhibition showed the strongest and most consistent effects among the three inhibitors across both cell lines, we next asked whether genetic targeting of *PAK1* also cooperates with RET TKI. CUTO32 and LC-2/Ad cells were treated with siRNA pools targeting *PAK1* and with pralsetinib. In CUTO32, the number of viable cells was not affected by *PAK1* knockdown alone, but *PAK1* silencing strongly synergized with pralsetinib treatment (Fig. 5B). A similar trend was observed in LC-2/Ad cells although it did not reach statistical significance (Supplementary Fig. S5). It is possible that these cells may additionally depend on other PAK family kinases like PAK2 or PAK4, which are also targets of PF-3758309 used in Fig. 5A. Furthermore, this could be also the result of weaker knockdown efficiency compared to CUTO32 cells (Fig. 5B and Supplementary Fig. 5). In summary, while AKT and mTORC1 inhibition may moderately contribute to the observed overall cellular synergy between RET TKIs and dasatinib, the major contribution was made by inhibition of PAK.

**Figure 5.**
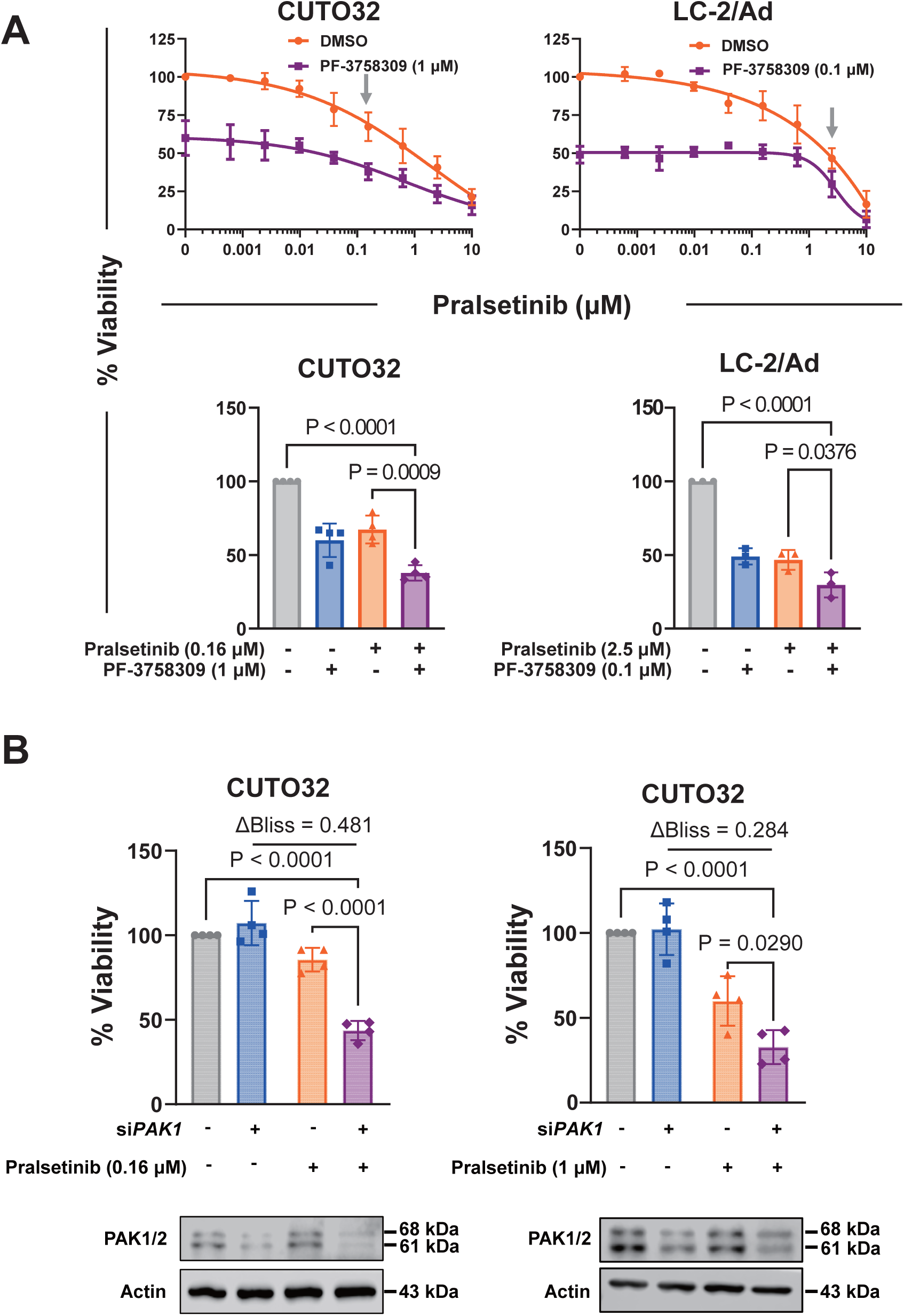
Effects of PAK targeting on modulation of pralsetinib sensitivity in *RET*^+^ NSCLC cells. (A) Viability of CUTO32 and LC-2/Ad cells upon co-treatment with pralsetinib and the PAK inhibitor PF-3758309 (CUTO32: 1 μM; LC2/Ad: 0.1 μM) for 3 days (Top). Cell viability of *RET*^+^ NSCLC cells at single concentrations of pralsetinib (CUTO32: 0.16 μM; LC-2/Ad: 2.5 μM, see grey arrows above) (Bottom). (B) Cell viability and Bliss synergy analysis of CUTO32 cells after siRNA-mediated *PAK1* knockdown (4 days) and pralsetinib treatment (0.16 and 1 μM) for 3 days (Top). PAK1 Western blot analysis of *PAK1* knockdown in CUTO32 cells (Bottom).

### 3.6. The next generation selective SRC inhibitor eCF506 synergizes with RET TKIs in *RET* fusion-positive cancer cells

Although dasatinib is an FDA-approved SRC TKI, its extensive off-target profile could cause unexpected adverse effects in clinical studies. In this regard, more selective SRC inhibitors may be more clinically beneficial options to combine with RET TKIs by minimizing off-target effects and associated toxicities. eCF506 (NXP900) is a recently developed SRC TKI, which inhibits both kinase activity and scaffolding functions of SRC by locking it in its inactive conformation [44]. This compound has high potency (IC_50_ less than 0.5 nM) comparable to dasatinib while displaying higher selectivity, and it is currently in early clinical trials for advanced cancers [44]. Thus, we determined whether eCF506 also exhibits synergy with RET TKIs in *RET*^+^ cancer cells. First, we analyzed signaling proteins, which we had observed to be downregulated by the combination of pralsetinib and dasatinib. This analysis showed that eCF506 also cooperated with pralsetinib to further inhibit the activity of PAK, AKT, and S6 in CUTO32 cells, similar to dasatinib (Fig. 6A). Next, we explored cell viability, clonogenicity and induction of apoptosis upon co-treatment with RET TKIs and eCF506 in *RET*^+^ NSCLC and PTC cell lines. In line with the observed signaling effects, eCF506 strongly synergized with RET TKIs to suppress viability and clonogenicity of *RET*^+^ cancer cells (Fig. 6B, C and Supplementary Fig. S6A-D). Finally, similar to dasatinib, addition of eCF506 strongly enhanced pralsetinib-induced apoptosis as indicated by increased PARP1 and caspase-3 cleavage (Fig. 6D). In summary, the selective next-generation clinical SRC TKI eCF506 strongly enhanced activity of RET TKIs in *RET*+ NSCLC.

**Figure 6.**
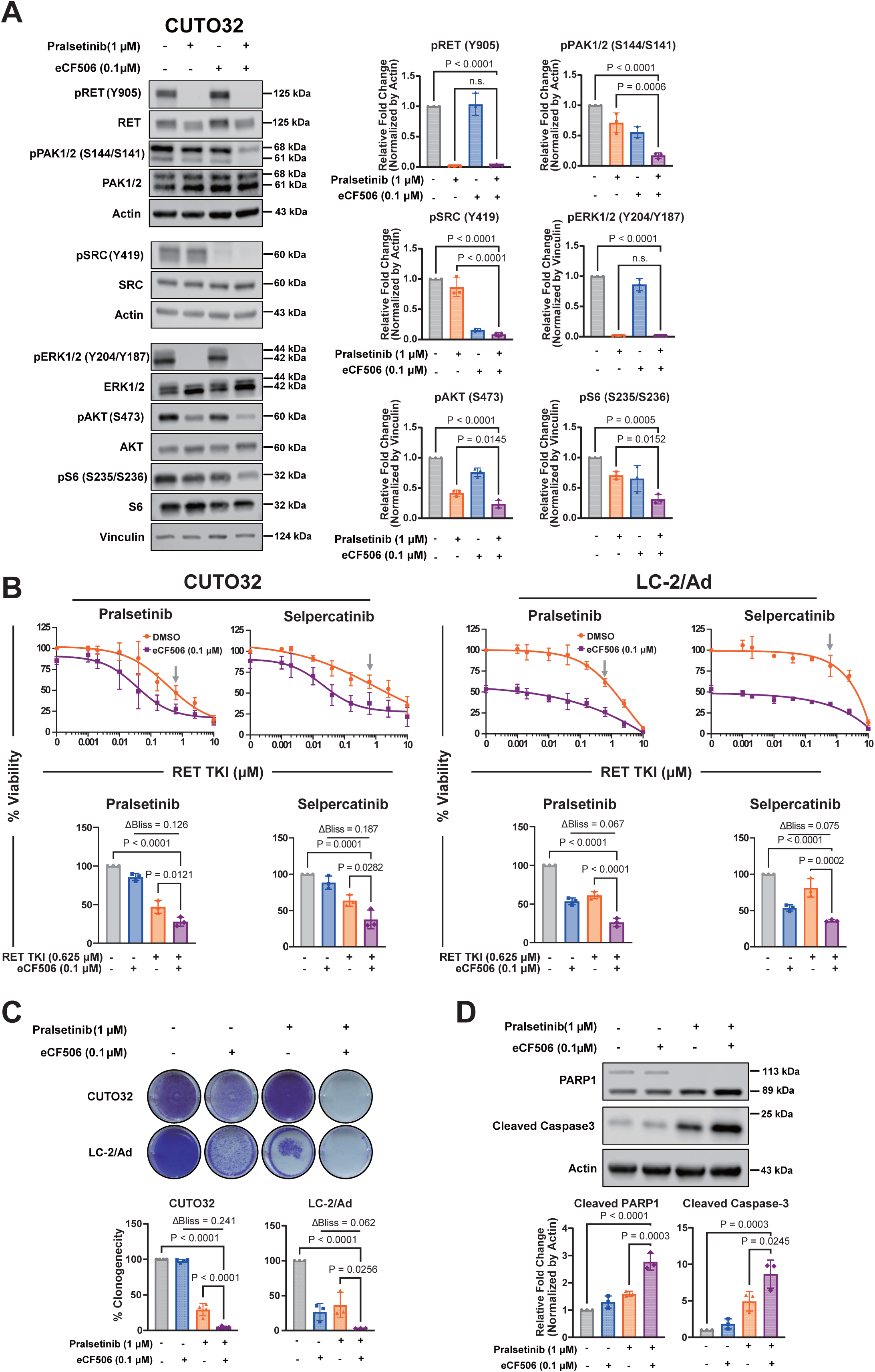
Combination effects of the next-generation SRC TKI eCF506 with pralsetinib in *RET*^+^ NSCLC cells. (A) (Left) Western blotting of indicated signaling proteins in CUTO32 cells after treatment with pralsetinib (1 μM), eCF506 (0.1 μM) or their combination for 3 hours. (Right) Quantification of p-RET, p-SRC, pPAK1/2, pERK1/2, pAKT, and p-S6 blots,normalized by loading controls (actin or vinculin). (B) (Top) Cell viability of CUTO32 and LC-2/Ad cells upon treatment with RET TKIs alone and in combination with eCF506 (0.1 μM) for 3 days. (Bottom) Cell viability and Bliss synergy analysis of *RET*^+^ NSCLC cells at single concentrations of RET TKIs (0.625 μM, see grey arrows above) and eCF506 (0.1 μM). (C) Clonogenic survival, quantification and Bliss synergy analysis of *RET*^+^ NSCLC cells after treatment with pralsetinib (1 μM), eCF506 (0.1 μM) or their combination for 7 days. (D) Western blotting and quantification of apoptosis markers upon treatment of CUTO32 cells with pralsetinib (1 μM), eCF506 (0.1 μM) and their combination for 3 days.

### 3.7. SRC inhibitors re-sensitize selpercatinib-resistant cells to RET inhibition

To determine whether combination strategies with dual inhibition of RET and SRC also have effects in *RET^+^* cells harboring acquired drug resistance, we generated a selpercatinib-resistant cell line following chronic exposure of *RET*^+^ TPC-1 cells to selpercatinib (2 μM) for 3 months. We then tested whether SRC inhibition was able to re-sensitize cells to selpercatinib. Viability and clonogenic assays showed that selpercatinib-resistant TPC-1 cells were much more resistant to single selperatinib treatment compared to the parental cells. Importantly, combination of selpercatinib and the SRC TKIs dasatinib or ecF506 were significantly more effective than the single agents to inhibit cell growth. (Fig. 7A, B). Taken together, these results suggest that SRC inhibition can re-sensitize TKI-resistant *RET*^+^ cells to selpercatinib.

**Figure 7.**
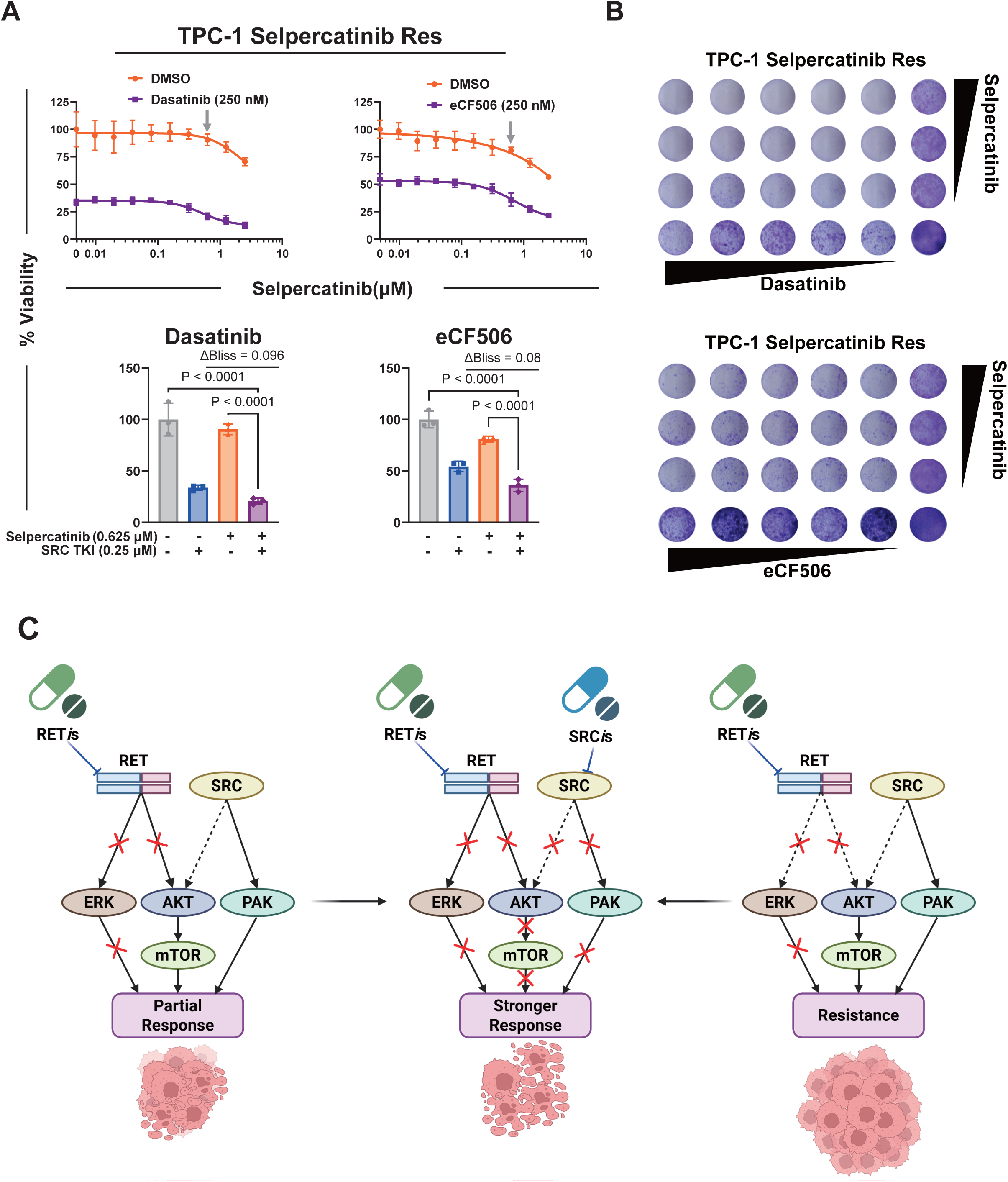
SRC inhibition resensitization of RET TKI-resistant *RET*^+^ PTC cells to selpercatinib. (A) Cell viability of selpercatinib-resistant TPC-1 cell line upon treatment with selpercatinib combined with dasatinib (0.25 µM) or eCF506 (0.25 µM) for 3 days (Top). Cell viability and Bliss synergy analysis of selpercatinib-resistant TPC-1 cells at single concentration of selpercatinib (0.625 μM, see grey arrows above) (Bottom). (B) Clonogenic survival of selpercatinib-resistant TPC-1 cells after selpercatinib treatment combined with dasatinib or eCF506 for 7 days. Cells were treated with 2-fold serial dilutions of each drug starting from 2.5 µM. (C) Schematic summary of molecular mechanisms through which SRC inhibition synergizes with RET TKIs in TKI-naïve (Left) and TKI-resistant (Right) *RET^+^* cancer cells.

## 4. Discussion

In this study, we found that SRC inhibitors sensitized *RET*^+^ NSCLC and PTC cells to current first-line selective RET TKIs (Fig. 7C). Rescue experiments with the drug-resistant SRC gatekeeper mutant furthermore validated that the drug combination effects were *SRC*-dependent as compared to through dasatinib off-targets. Phosphoproteomics analysis and subsequent functional validation identified that these synergistic effects were mediated by improved inhibition of PAK and AKT/mTORC1 signaling pathways downstream of SRC. Lastly, there may be an opportunity to use the next-generation clinical SRC inhibitor, eCF506, which has similar effects and higher SRC selectivity compared to dasatinib, as an alternative drug for clinical translation in combination with currently approved RET TKIs. While we focused on *RET*-independent signaling mechanisms in this study, this research does not include any on-target resistance mechanisms, which can disrupt the activity of pralsetinib and selpercatinib. Both drugs are vulnerable to secondary RET solvent front and gatekeeper mutations which sterically hinder the drug binding of RET due to substitution of valine at position 804 or glycine at position 810 with bulky, charged or polar residues [12]. In this case, one may consider treating *RET*^+^ cancers with the novel RET TKI, vepafestinib, which has activity against secondary RET mutants, and combining it with SRC inhibitors, such as eCF506 [13].

Although previous studies from different groups have suggested the association between RET and SRC, the study here provides a unique insight into the biology of *RET*^+^ cancers and outlines new opportunities for clinical translation. Kang *et al.*, reported the role of SRC in acquired resistance to the multikinase inhibitor dovitinib, which inhibits RET in LC-2/Ad cells [25]. However, dovitinib is not an approved drug for *RET*^+^ NSCLC and it has a large number of targets, including VEGFRs, PDGFRs and FGFRs, which drastically increases the risk for toxic off-target effects [45]. Another study demonstrated the vulnerability of PTC cells to SRC TKIs; however, combination with RET TKIs was not investigated as these drugs were not yet available at that time [26]. Thus, further cellular and mechanistic evidence was needed to validate the functions of SRC in *RET*^+^ cancer cells in the context of pralsetinib and selpercatinib, which are current frontline therapies. Our results illustrate that *RET^+^* NSCLC and PTC cells, which were not completely responsive to these RET TKIs, became more sensitive when SRC inhibitors were added. Furthermore, this study is significant as the combination treatment was effective in parental *RET*^+^ cancer cells, as well as PTC cells with acquired RET TKI resistance. While we identified SRC as a major protein accounting for bypass signaling mechanisms, alterations of other genes such as *KEAP1* mutation, amplification of *MET*, *KRAS* or *CDK2N2A* have also been reported [10]. This implies that TKI resistance mechanisms can be context-dependent and driven by proteins other than SRC, possibly necessitating adjustment of the therapeutic strategy in future research. For example, CUTO32 cells, which have *KIF5B-RET* (K15;R12) with *NF1* and *TP53* mutations were also sensitive to PLK1 and AURKA inhibitors, while CUTO22 cells expressing *KIF5B-RET* (K23;R12) with *NF1* and *APC* mutations were not responsive to those drugs [28]. Considering the heterogeneity of *RET*-dependent and *RET*-independent acquired resistance mechanisms it may be most beneficial to apply combinations of RET and SRC TKIs in the treatment-naïve setting.

The present study identified PAKs as major SRC downstream target proteins, which mediate much of the bypass signaling mechanisms, in *RET*^+^ NSCLC. Besides their role in cell survival and proliferation, PAKs regulate downstream effectors including LIMK1 and αvβ5 integrin to reorganize cytoskeleton structures, which are intimately related with cell motility, invasion, and eventually metastases [40]. In addition, PAKs have been shown to inhibit Hippo signaling leading to downstream activation of the transcriptional coactivator YAP1, dysregulation of which is highly correlated with cancer progression [46]. This is consistent with a recent study reporting that *RET^+^* cancer cells can acquire resistance to RET TKIs via YAP1-mediated activation of ERBB3 signaling [47]. Thus, PAK family kinases may be promising co-targets, alternatively to SRC, for *RET*^+^ NSCLCs, which are often diagnosed at advanced stages with established metastases [8, 9]. However, no PAK inhibitors are currently clinically available. A phase I clinical trial using PF-3758309 to treat solid tumors was terminated early due to unexpected side effects, while another phase I clinical trial using KPT-9274, which targets PAK4 and NAMPT, is in progress [48]. Another option may be using the clinical-stage compound MBQ-167, which selectively inhibits RAC1/CDC42 upstream of PAK [49]. More studies comparing the extent of drug efficacy shared between RAC1/PAK and SRC inhibitors would be required.

## 5. Conclusion

In conclusion, this study has revealed synergistic effects of RET and SRC TKIs in *RET*^+^ cancer cells. The molecular mechanisms underlying these effects are mediated by on-target SRC inhibition and suppression of downstream PAK and AKT/mTOR signaling. For improvements of clinical outcomes to treat patients with *RET^+^* cancers, the next-generation SRC TKI eCF506 or, alternatively, RAC/PAK inhibitors may be promising options as these combination treatments, particularly in the upfront setting, may enhance the clinical efficacy of RET-targeted therapies, thus overcoming or delaying *RET*-independent acquired drug resistance mechanisms.

## Supporting information

Combined supplementary figures

## Acknowledgements

This work has been supported by the Department of Defense Lung Cancer Research Program W81XWH-22-1-0607 (to U.R.), the H. Lee Moffitt Cancer Center & Research Institute, the Moffitt Lung Cancer Center of Excellence. E.C. gratefully acknowledges support from the Lung Cancer Research Foundation (LCRF), the Madelon Ravlin Grant Memorial Award from the Woman’s Cancer Association of the University of Miami, the Sylvester ACS Pilot Project, the 2025 Catalyst award from the Sylvester Comprehensive Cancer Center (SCCC), the Dwoskin family donation, and the Tumor biology intra-programmatic pilot award from the SCCC. E.C. also thanks the LCRF for the 2022 William C. Rippe Award. A.D. was supported by the National Cancer Institute/National Institutes of Health P30-CA008748. We also wish to acknowledge the Proteomics and Metabolomics Core, Molecular Genomics Core, and Biostatistics and Bioinformatics Shared Resource at the H. Lee Moffitt Cancer Center & Research Institute; Moffitt core facilities are supported by the NCI Cancer Center Support grant (P30-CA076292). We would also like to thank Dr. Robert Doebele for providing CUTO32 and LC-2/Ad cell lines.

## Author contributions

Conception and design: J.S., J.M.K., E.C., and U.R. Development of methodology: J.S., L.L.R.R. Acquisition of data: J.S., B.F., N.V.B., V.F., P.R., and N.S.-C. Analysis and interpretation of data: J.S., L.L.R.R., B.F., and E.A.W. Administrative, technical, or material support: E.B.H., and A.D. Writing of the manuscript: J.S., E.C. and U.R. Study supervision: E.C. and U.R. All authors read, edited, and approved the final manuscript.

## Data accessibility

All data reported in this paper will be shared by the lead contacts upon request. The mass spectrometry proteomics data have been deposited to the ProteomeXchange Consortium via the PRIDE database (http://www.proteomexchange.org) with the dataset identifier PXD063113 and 10.6019/PXD063113, and are publicly available as of the date of publication.

**Supplementary** Figure 1. Combination effects of RET TKIs with dasatinib in *RET*^+^ TPC-1 PTC cells. (A) Cell viability of *RET*^+^ TPC-1 PTC cells upon treatment with RET TKIs (pralsetinib, selpercatinib) and dasatinib (0.25 μM) for 3 days (Top). Cell viability and Bliss synergy analysis of TPC-1 cells at single concentration of RET TKIs (15 nM) (Bottom). (B) Clonogenic survival of TPC-1 cells after treatment with RET TKIs and dasatinib for 7 days. Cells were treated with 2-fold serial dilutions of each drug starting from 2.5 µM.

**Supplementary** Figure 2. SRC-dependency of synergistic effects between RET TKIs and dasatinib in LC-2/Ad *RET*^+^ NSCLC cells. (A) SRC autophosphorylation in LC-2/Ad cells expressing SRC wild-type (WT) or SRC T341I gatekeeper mutation upon treatment with dasatinib (0.1 μM) for 3 hours. (B) Cell viability of LC-2/Ad cells expressing SRC WT or SRC T341I gatekeeper mutation after combined treatment with dasatinib (0.16 μM) and RET TKIs pralsetinib and selpercatinib (both 1 μM) for 3 days. (C) Clonogenic survival and quantification of LC-2/Ad expressing SRC WT or SRC T341I gatekeeper mutation upon treatment with pralsetinib (1 μM), dasatinib (0.1 μM) or combination thereof for 7 days.

**Supplementary** Figure 3. Protein-protein network analysis of proteomics data in *RET^+^* CUTO32 NSCLC cells. (A) STRING-based protein-protein network analysis and GLay-based subclustering of *RET*^+^ CUTO32 NSCLC cells after co-treatment with pralsetinib (1 μM) and dasatinib (0.1 μM) for 3 hours. All regulated proteins from phosphoproteomics and expression proteomics data were combined. (B) Scatter plot based on fold changes (phosphorylation or total expression) and betweenness centrality of proteins significantly regulated by combination treatment. Color gradient represents log_2_ fold change of signals.

**Supplementary** Figure 4. Effects of selective inhibitions of AKT, mTOR and PAK. (A) (Top) Cell viability of *RET*^+^ NSCLC cells upon treatment with pralsetinib combined with the AKT inhibitor GSK690693 (1 μM) or the mTOR inhibitor rapamycin (10 nM). (Bottom) Cell viability at the single concentration of RET TKIs (CUTO32: 0.16 μM; LC-2/Ad: 2.5 μM). (B) Phosphorylation of PAK1/2, AKT and S6 in CUTO32 cells after treatment with PF-3758309 at indicated concentrations for 3 hours.

**Supplementary** Figure 5. Silencing of *PAK1* in *RET^+^* LC2/Ad NSCLC cells. (Top) Cell viability of LC-2/Ad cells after siRNA-mediated *PAK1* knockdown (4 days) and pralsetinib treatment (2.5 μM, 3 days). (Bottom) Western blotting of *PAK1* knockdown in LC-2/ad cells.

**Supplementary** Figure 6. Combination effects of RET TKIs with eCF506 in *RET*^+^ TPC-1 PTC cells. (A) (Top) Cell viability of TPC-1 cells upon treatment with RET TKIs alone and combined with eCF506 (0.25 μM) for 3 days. (Bottom) Cell viability and Bliss synergy analysis of TPC-1 cells at single concentration of RET TKIs (15 nM). (B) Clonogenic survival of TPC-1 cells after treatment with RET TKIs, eCF506 or their combination for 7 days. Cells were treated with 2-fold serial dilutions of each drug starting from 2.5 µM.

**Supplementary Table S1.** Global phosphoproteomics (pSTY) data, related to Figure 3. MaxQuant-analyzed data for pSTY phosphopeptides detected in CUTO32 cells treated with pralsetinib (1 μM), dasatinib (0.1 μM) or combination thereof for 3 hours.

**Supplementary Table S2.** Expression proteomics data, related to Figure 3. MaxQuant-analyzed data for total protein expression detected in CUTO32 cells treated with pralsetinib (1 μM), dasatinib (0.1 μM) or combination thereof for 3 hours.

