## Supplementary material for "Dual targeting of RET and SRC synergizes in RET fusion-positive cancer cells": Combined supplementary figures

### Supplementary Fig. S1.

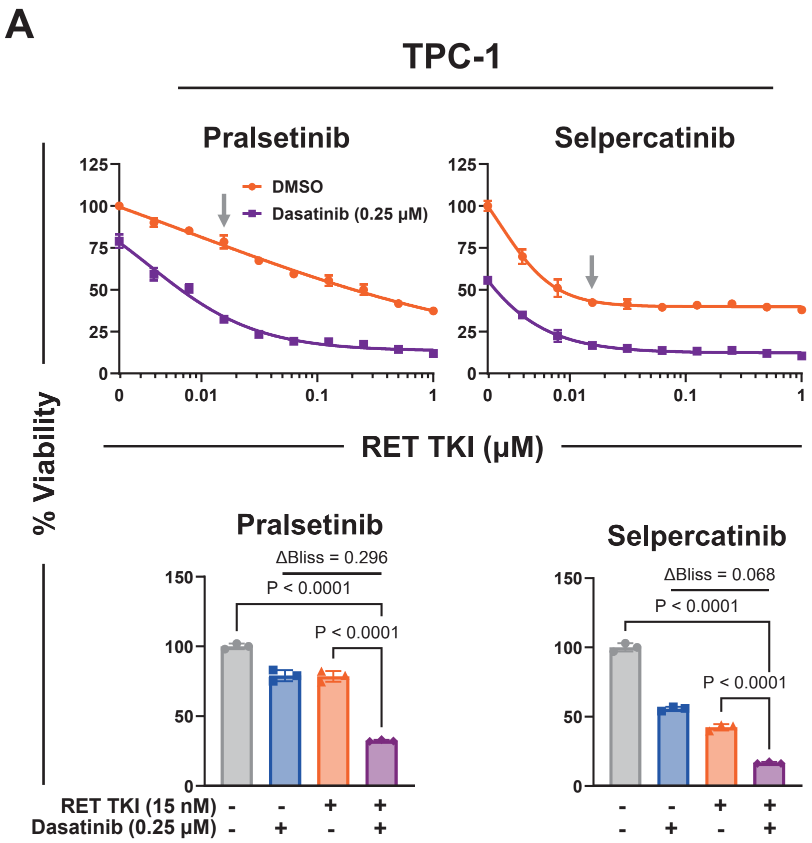


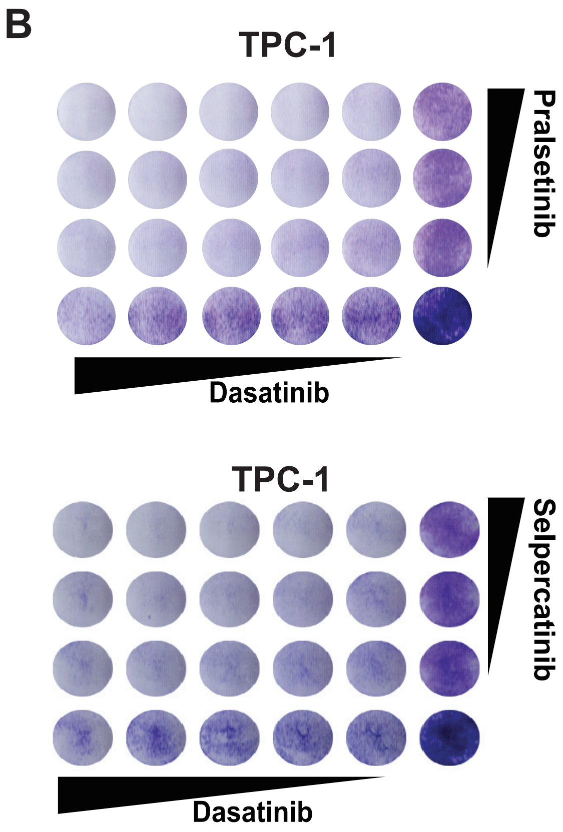

### Supplementary Fig. S2.

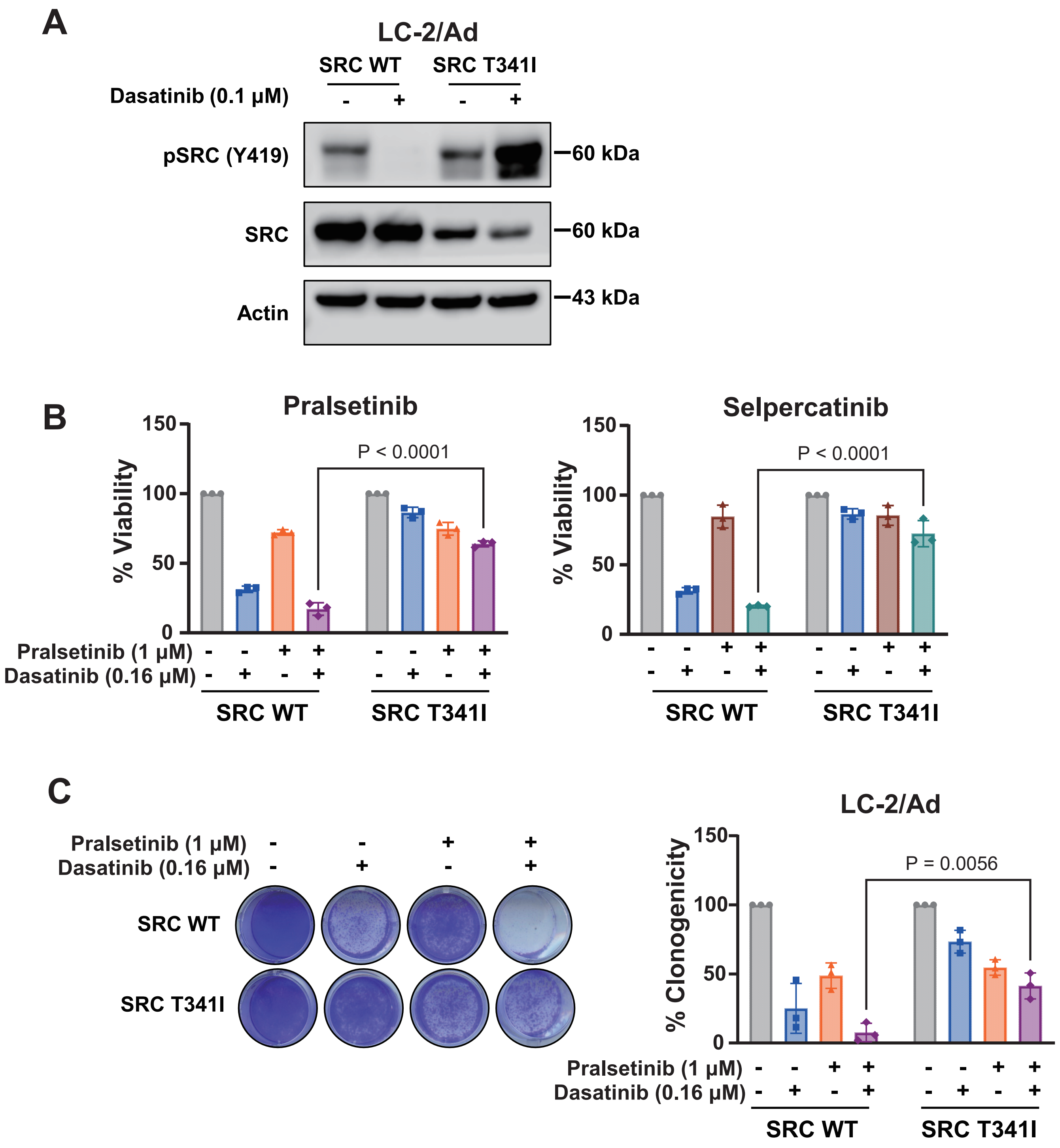

### Supplementary Fig. S3

**A**

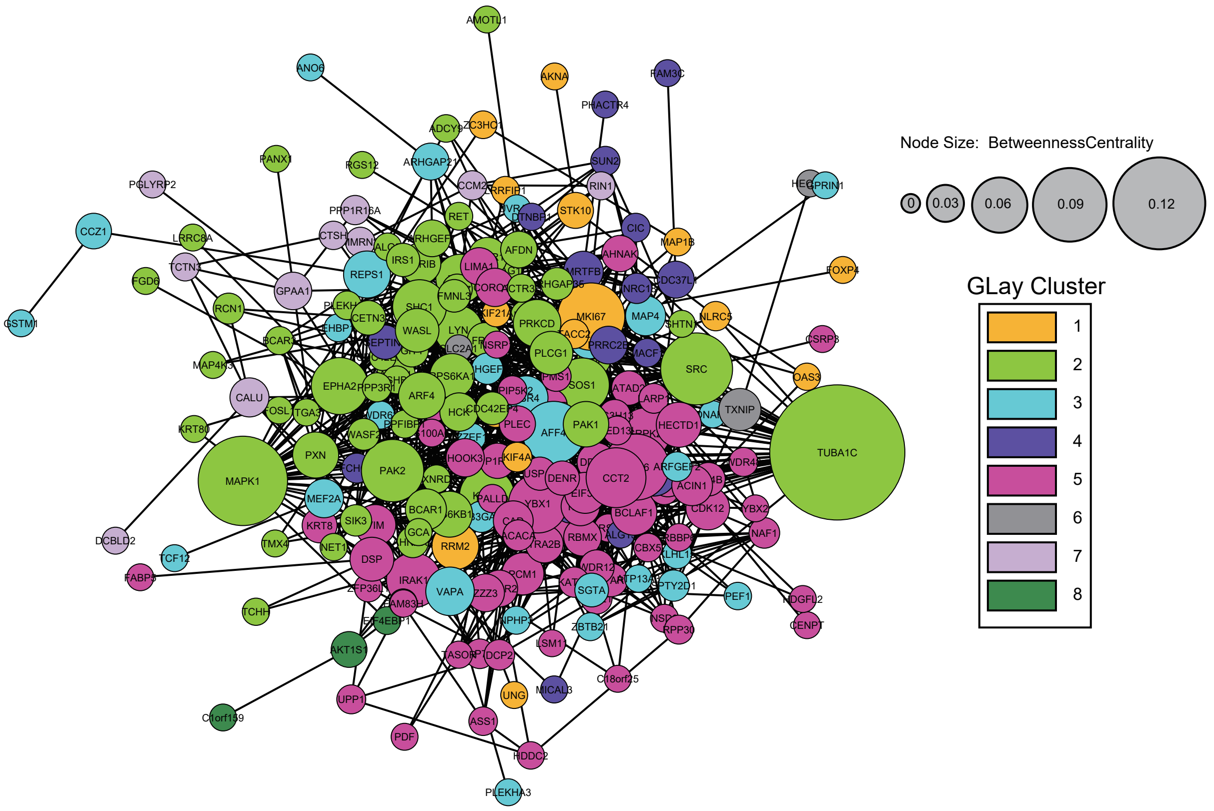


**B**

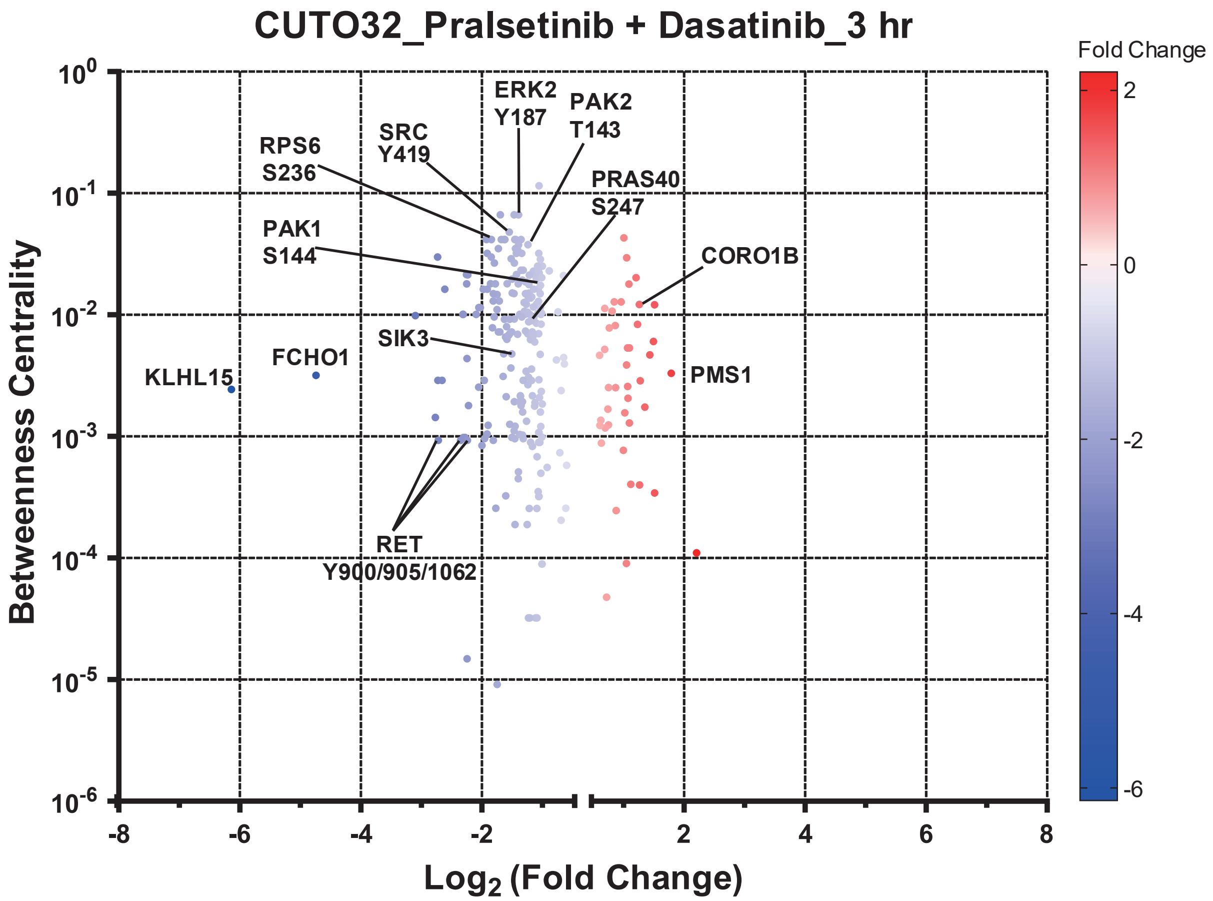

### Supplementary Fig. S4

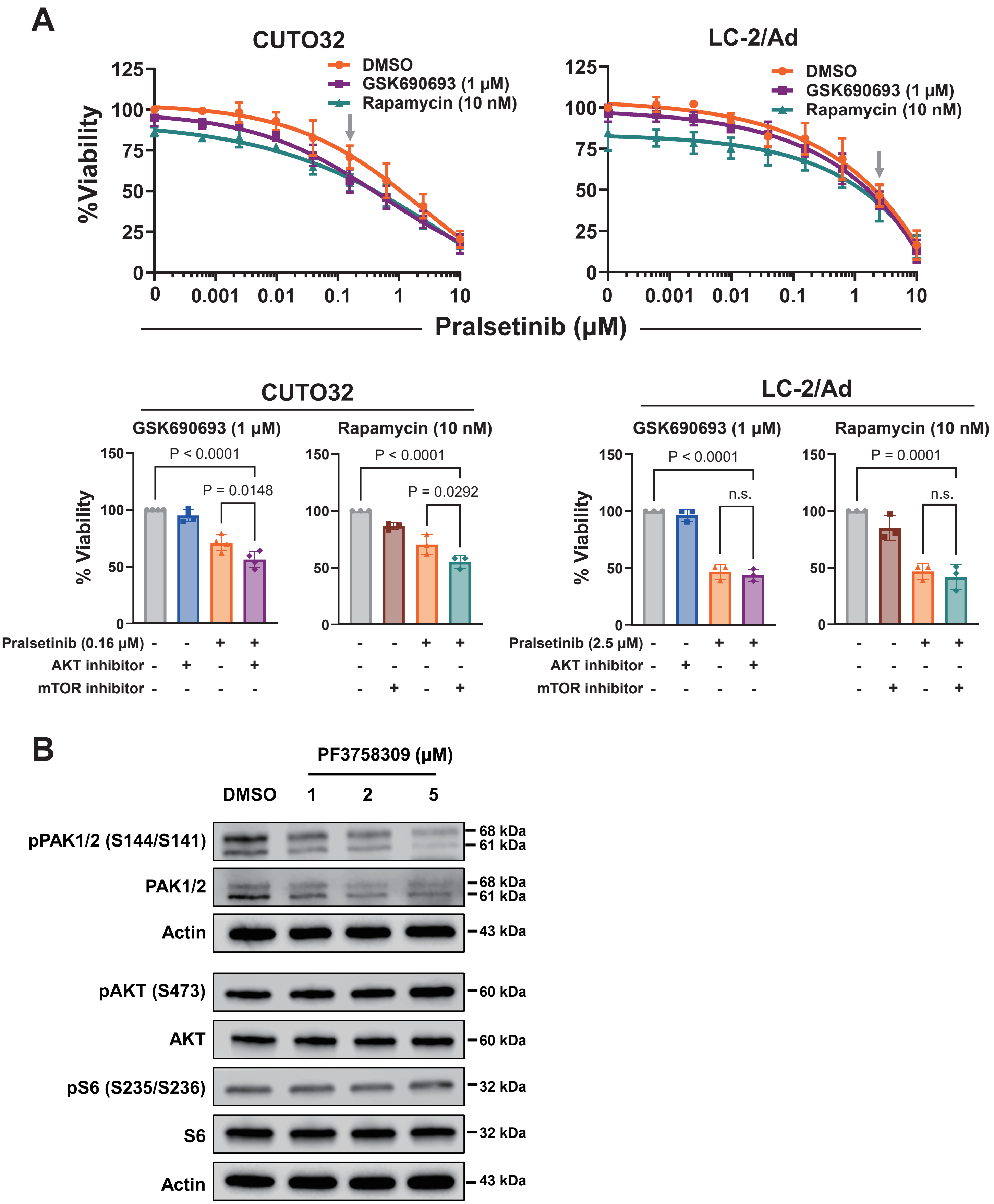

**Supplementary Fig. S5**

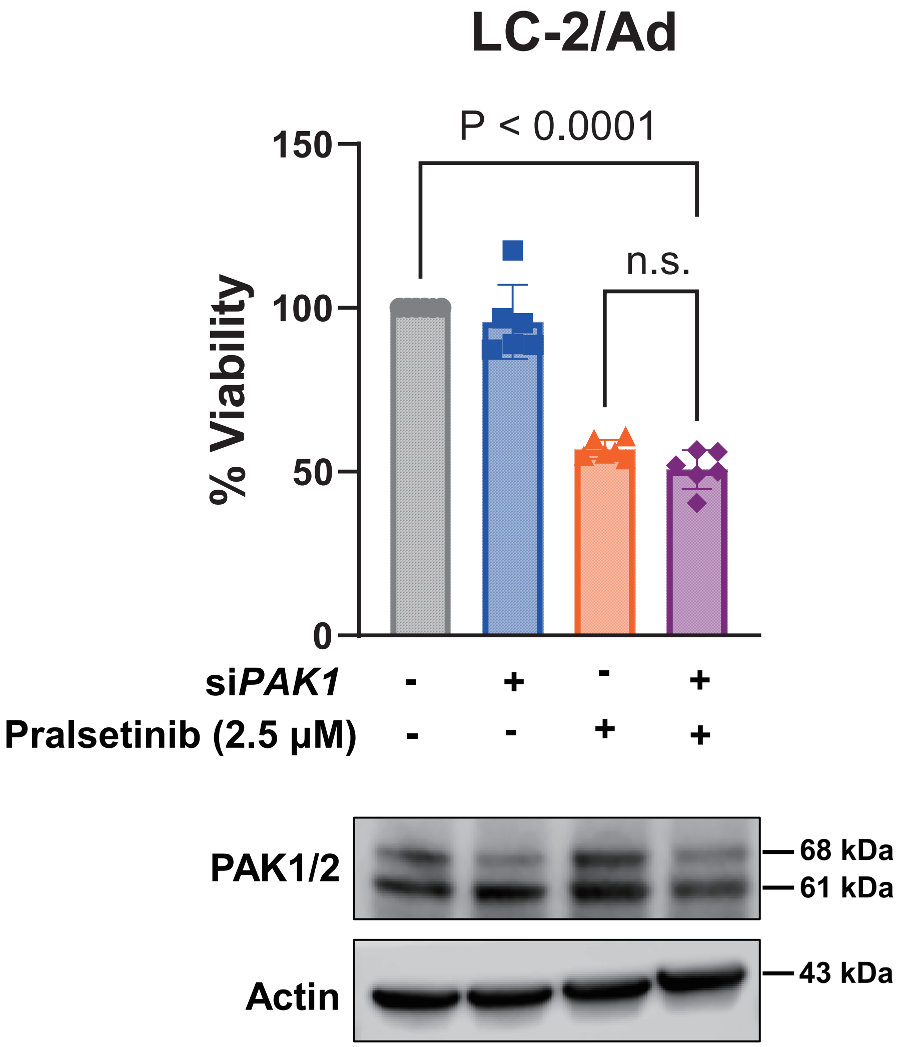

### Supplementary Fig. S6.

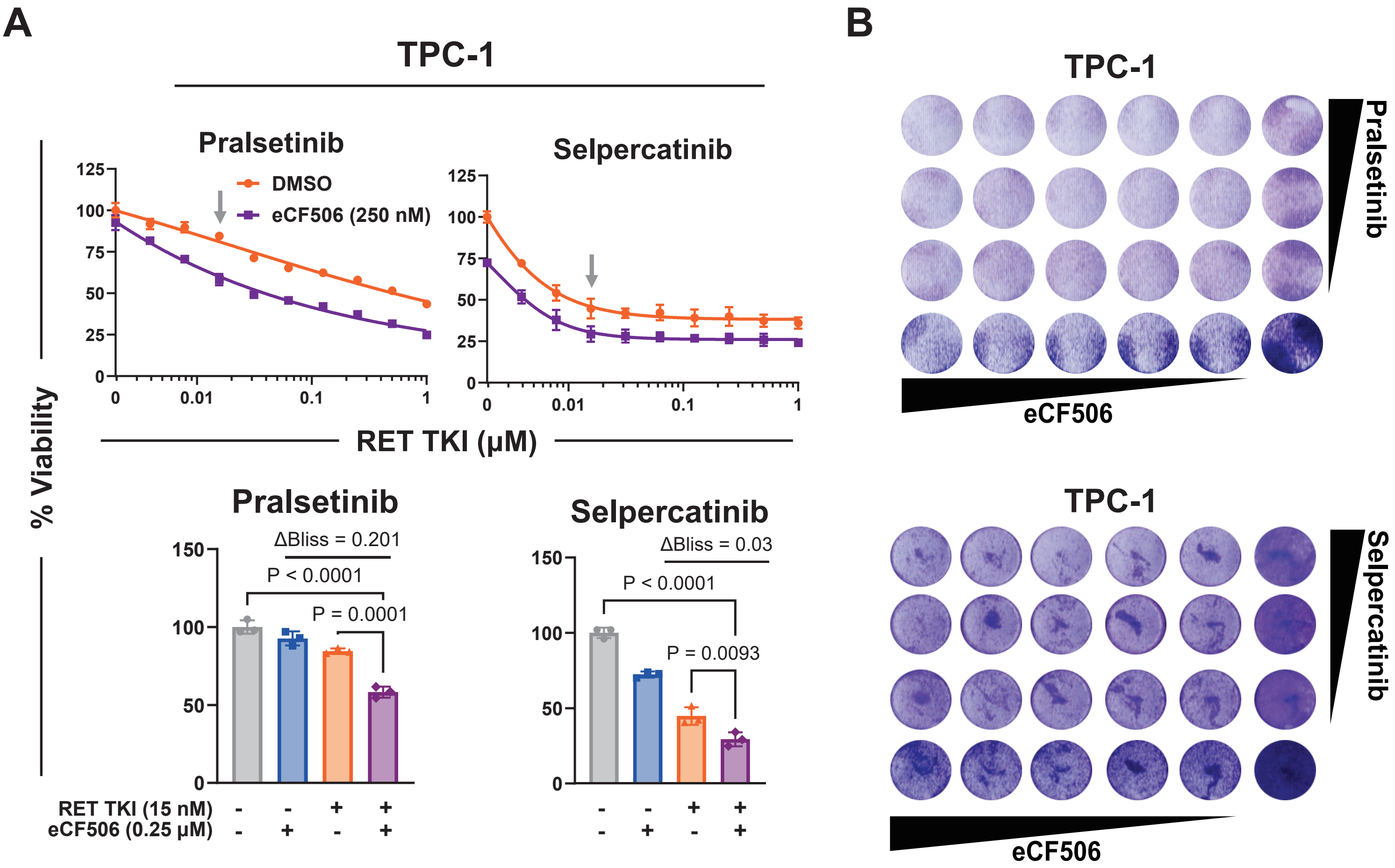
